# Temperature-Jump Solution X-ray Scattering Reveals Distinct Motions in a Dynamic Enzyme

**DOI:** 10.1101/476432

**Authors:** Michael C. Thompson, Benjamin A. Barad, Alexander M. Wolff, Hyun Sun Cho, Friedrich Schotte, Daniel M.C. Schwarz, Philip Anfinrud, James S. Fraser

## Abstract

Correlated motions of proteins and their bound solvent molecules are critical to function, but these features are difficult to resolve using traditional structure determination techniques. Time-resolved methods hold promise for addressing this challenge but have relied on the exploitation of exotic protein photoactivity, and are therefore not generalizable. Temperature-jumps (T-jumps), through thermal excitation of the solvent, have been implemented to study protein dynamics using spectroscopic techniques, but their implementation in X-ray scattering experiments has been limited. Here, we perform T-jump small- and wide-angle X-ray scattering (SAXS/WAXS) measurements on a dynamic enzyme, cyclophilin A (CypA), demonstrating that these experiments are able to capture functional intramolecular protein dynamics. We show that CypA displays rich dynamics following a T-jump, and use the resulting time-resolved signal to assess the kinetics of conformational changes in the enzyme. Two relaxation processes are resolved, which can be characterized by Arrhenius behavior. We also used mutations that have distinct functional effects to disentangle the relationship of the observed relaxation processes. A fast process is related to surface loop motions important for substrate specificity, whereas a slower process is related to motions in the core of the protein that are critical for catalytic turnover. These results demonstrate the power of time-resolved X-ray scattering experiments for characterizing protein and solvent dynamics on the μs-ms timescale. We expect the T-jump methodology presented here will be useful for understanding kinetic correlations between local conformational changes of proteins and their bound solvent molecules, which are poorly explained by the results of traditional, static measurements of molecular structure.

## INTRODUCTION

Protein motions are critical for functions such as enzyme catalysis and allosteric signal transduction (Henzler-Wildman and Kern, 2007), but it remains challenging to study excursions away from the most populated conformations (van den Bedem and Fraser, 2015). Traditional methods that utilize X-rays for structural characterization of biological macromolecules, such as crystallography and solution scattering, provide high-quality structural information, but this information is both spatially and temporally averaged because the measurements are performed on large ensembles of molecules and are typically slower than the timescales of molecular motion (van den Bedem and Fraser, 2015; Bottaro and Lindorff-Larsen, 2018). To some extent, the spatial averaging inherent to X-ray experiments is advantageous, because it reveals the alternative local conformations of a molecule that are significantly populated at equilibrium; however, structural states that are not significantly populated at equilibrium, such as intermediates along a conformational transition pathway, are effectively invisible. The temporal averaging inherent to X-ray experiments also results in a loss of information about how transitions between local alternative conformational states are coupled to one another. To gain kinetic information about molecular motion, researchers often turn to spectroscopic methods, but it can be difficult to correlate spectroscopic observables with high resolution structural models.

Time-resolved X-ray scattering and diffraction can overcome the limitations of traditional structure determination for studying the dynamics of biomolecules (Cho et al., 2016; Neutze and Moffat, 2012; Schmidt, 2017; Schotte et al., 2012). In these experiments, a fast perturbation is applied to the sample to remove it from conformational equilibrium and synchronize conformational changes in a significant fraction of the molecules. Ultrafast X-ray pulses, which are short relative to motions of interest, are then used to perform structural measurements in real time as the system relaxes to a new equilibrium, providing simultaneous structural and kinetic information at high spatial and temporal resolution. Time-resolved X-ray experiments can identify transiently-populated structural states along a conformational transition pathway, and reveal kinetic couplings between conformations (Schlichting and Miao, 2012). Despite this potential to provide a wealth of information, especially when combined with molecular dynamics simulation (Arnlund et al., 2014; Berntsson et al., 2017; Brinkmann and Hub, 2016; Takala et al., 2014), time-resolved experiments have not been broadly applied by structural biologists. To date, systems that have been most rigorously studied are those in which a protein conformational change is coupled to excitation of a photoactive ligand molecule, because the conformational change can be initiated with an ultrafast optical laser pulse (e.g. (Barends et al., 2015; Coquelle et al., 2018; Kern et al., 2018; Malmerberg et al., 2011; Nogly et al., 2018; Pande et al., 2016)). Unfortunately, the number of proteins that undergo specific photochemistry as part of their functional cycle is small, and there is a fundamental need to develop generalized methods that can be used to synchronously excite conformational transitions in any protein molecule and expand the utility of time-resolved structural experiments (Hekstra et al., 2016).

**Figure 1.**
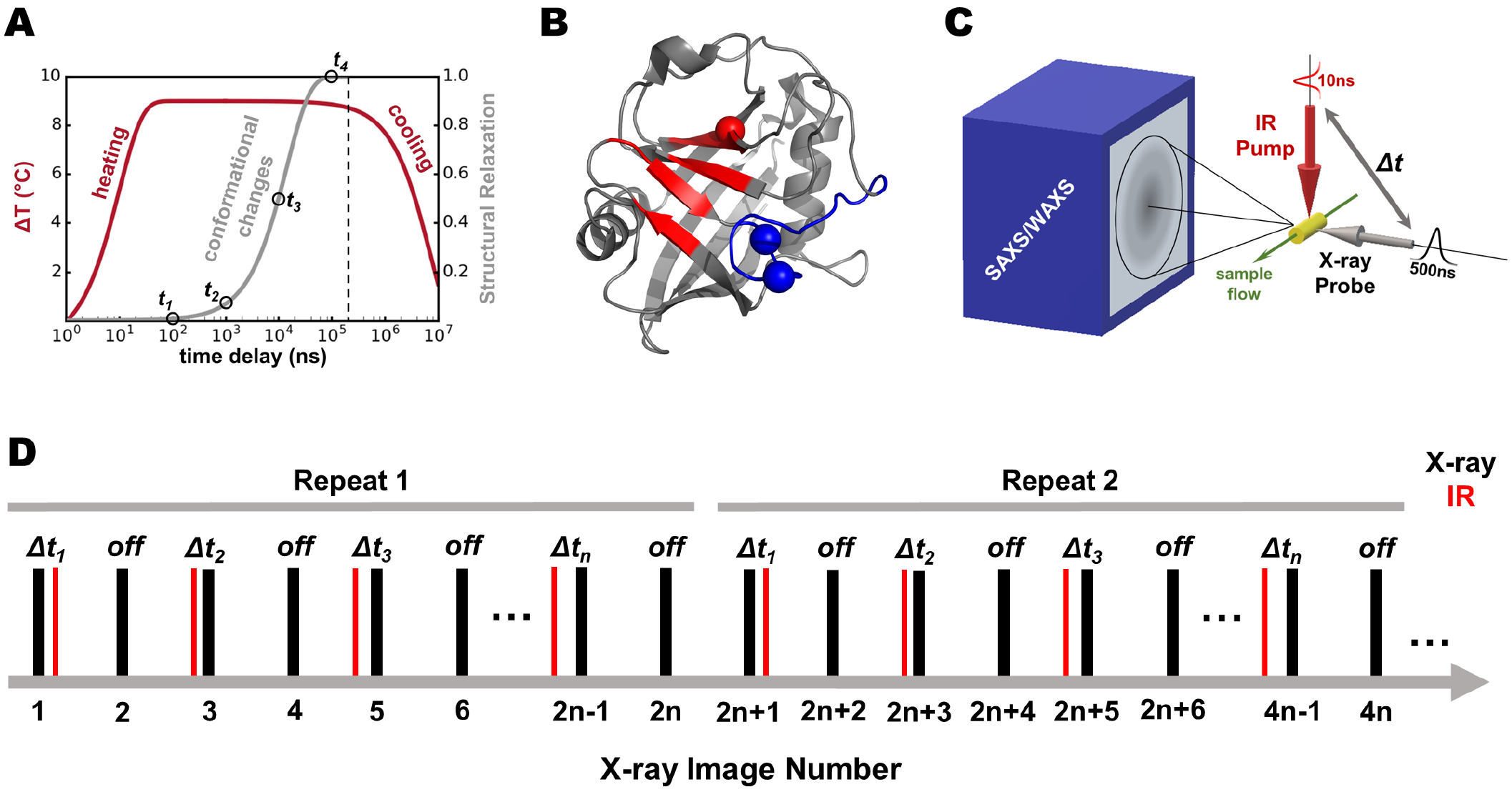
Overview of T-jump SAXS/WAXS experiments. A.) During a T-jump experiment, an infrared (IR) laser pulse, several nanoseconds in duration, vibrationally excites the water O-H stretch and rapidly heats an aqueous solution of protein molecules (red curve). Although heating is fast, the kinetic barriers to protein motions cause a lag in the structural relaxation to a new thermal equilibrium of conformational states (gray curve). B) Ribbon diagram depicting a single cyclophilin A (CypA) molecule. The “core” dynamic residues that are linked to catalysis are colored red, and the site of a key mutation (S99T) is identified by a sphere at its Cα position. Likewise, the “loop” region adjacent to the active site that helps determine substrate specificity is colored blue, and the site of key mutations (D66N/R69H) are also identified by spheres at their Cα positions. C) A schematic depicting the T-jump SAXS/WAXS instrumentation is shown with key features highlighted. A liquid sample flows horizontally through the interaction region, where it interacts with mutually perpendicular IR pump and X-ray probe beams. Both the pump and probe sources are pulsed, with a defined time delay between their arrival at the sample. Small- and wide-angle X-ray scattering (SAXS/WAXS) patterns are recorded on a single detector panel. D) The diagram illustrates the data collection sequence used for the experiments described here. For each pump-probe time delay, a pair of images was collected such that the first image was a pump-probe measurement (“laser on”) and the subsequent image was collected with no application of the pump laser (“laser off”). On-off pairs with increasing pump-probe time delays were measured in succession until all of the desired delay times were acquired, and this sequence was repeated as many as 50 times to improve the signal-to-noise ratio of the data. Note that the first measurement within each repeat is a control measurement, wherein the probe pulse arrived at the sample before the pump pulse (negative time delay).

Protein structural dynamics are intimately coupled to the thermal fluctuation of the surrounding solvent (“solvent slaving” (Fenimore et al., 2002; Frauenfelder et al., 2007)), and thermal excitation of the solvent by infrared (IR) laser temperature-jump has been used in numerous pump-probe experiments. These experiments work on the principle that absorption of IR photons excites the O-H stretching modes of water molecules, and the increased vibrational energy is dissipated through increased rotation and translation of the solvent molecules, effectively converting electromagnetic energy into kinetic (thermal) energy. Because this process of solvent heating and subsequent heat transfer to the protein is much faster than the large-scale molecular motions that define protein conformational changes, the sudden T-jump removes conformational ensembles of protein molecules from their thermal equilibrium so that their structural dynamics can be measured using relaxation methods (Figure 1A). For example, T-jump perturbations have been coupled to ultrafast spectroscopic methods, including Fourier-transform infrared (FTIR) spectroscopy (Wang and El-Sayed, 1999; Wang et al., 2005), nuclear magnetic resonance (NMR) (Akasaka et al., 1991; Gillespie et al., 2003; Yamasaki et al., 2013), and various forms of fluorescence spectroscopy (Meadows et al., 2015; Vaughn et al., 2018), for the study of protein folding and enzyme dynamics. While these methods provide detailed kinetic information, they yield only very limited structural information about the underlying atomic ensemble. In contrast, the application of T-jumps to time-resolved X-ray scattering and diffraction has been very limited. Nearly two decades ago, Hori, et al used temperature-jump Laue crystallography to study the initial unfolding step of 3-isopropylmalate dehydrogenase (Hori et al., 2000). That study explored only a single pump-probe time delay, which allowed them to observe laser-induced structural changes but precluded kinetic analysis. Within the last two years, the laser T-jump method has been paired with X-ray solution scattering to explore the oligomerization of insulin in non-physiological conditions (Rimmerman et al., 2017, 2018) and hemoglobin (Cho et al., 2018). The results and analysis we present here expand the role of the T-jump method in structural biology, by demonstrating that T-jump X-ray scattering experiments can be used as a general method to explore the functional, internal dynamics of proteins under solution conditions. Additionally, we provide a detailed outline of a data reduction and analysis procedure suitable for T-jump SAXS/WAXS experiments.

The T-jump SAXS/WAXS experiments we describe here used human cyclophilin A (CypA), a proline isomerase enzyme that functions as a protein folding chaperone and as a modulator of intracellular signaling pathways. CypA has been the subject of many NMR experiments that have identified two primary dynamic features of interest (Figure 1B). First, the active site-adjacent loops (covering approximately residues 60-80 and hereafter referred to as the “loops” region)) are mobile on a ms-timescale (Eisenmesser et al., 2005). This region is especially interesting because evolutionarily selected mutations along these loops perturb the dynamics of the loop (Caines et al., 2012), alter the binding specificity of CypA for substrates such as HIV capsids (Price et al., 2009), and restrict the host range of these viruses (Virgen et al., 2008; Wilson et al., 2008). Second, a group of residues that extends from the active site into the core of the protein has also been shown to be mobile on a ms-timescale (Eisenmesser et al., 2005). Subsequent work incorporating multi-temperature X-ray crystallography (Keedy et al., 2015), mutagenesis (Fraser et al., 2009), and further NMR experiments (Otten et al., 2018) have established a relationship between the conformational dynamics of a group of side chains in this region and catalysis. Motivated by the sensitivity of the conformational state of the active site-core network (hereafter referred to as the “core” region) to temperature (Keedy et al., 2015), we performed infrared laser-driven T-jumps on buffered aqueous solutions of CypA and measured subsequent, time-dependent changes in small and wide angle X-ray scattering (SAXS/ WAXS). While our measurements provide only low resolution structural information, we were able to measure the kinetics of protein conformational changes in CypA. We identified two relaxation processes, and by performing T-jump experiments at a range of different temperatures, we were able to calculate thermodynamic properties of the transition states for the underlying conformational transitions. Specific mutants in the “loops” or the “core” regions of CypA show that the two processes are independent, each representing a distinct and uncoupled reaction coordinate on a complex conformational landscape. Collectively, our measurements and analysis demonstrate that a wealth of information about a protein’s conformational landscape can be obtained by pairing laser-induced T-jump with time-resolved X-ray scattering.

## RESULTS

### A method for simultaneous measurement of structural and kinetic details of intrinsic protein dynamics

To measure protein structural dynamics, we utilized a pump-probe method that pairs an infrared laser-induced temperature-jump with global measurement of protein structure via X-ray solution scattering (Figure 1C). We performed solvent heating in aqueous protein solutions by exciting the water O-H stretch with mid-IR laser pulses (1443nm, 7ns duration). At regularly defined time delays following the IR heating pulse (from 562ns to 1ms), we probed the sample with high-brilliance synchrotron X-ray pulses from a pink-beam undulator (3% bandwidth at 12keV, **Supplemental Figure 1**) that were approximately 500ns in duration, and measured X-ray scattering using a large CCD detector that was capable of capturing small and wide scattering angles on a single panel. Because the duration of the IR pump pulse was sufficiently short compared the the duration of the X-ray probe, the heating was effectively instantaneous with respect to the relaxation processes we were able to observe. Data were collected as interleaved “laser on” and “laser off” X-ray scattering images, so that each pump-probe measurement could be paired to a measurement made immediately before application of the pump laser (Figure 1D). We measured 27 unique pump-probe time delays across four decades of time spanning from 562ns to 1ms, performing 50 repeat measurements for each time delay. For each detector image, the individual pixel values were azimuthally averaged as a function of the scattering vector magnitude, *q*, to give one-dimensional scattering intensity profiles (*I(q)* curves). All scattering profiles were scaled to a single reference, and the data were analyzed as described below. These pump-probe measurements allowed us to monitor structural changes within the ensemble of heated molecules in real time as the system relaxed to a new thermal equilibrium following T-jump (Figure 1A).

### Calibrating the magnitude of the Temperature-Jump by Singular Value Decomposition

Because the isothermal compressibility of liquid water is highly temperature-dependent (Clark et al., 2010), X-ray scattering from the bulk solvent acts as an exquisitely sensitive thermometer that can be used to calibrate the magnitude of the T-jump in our experiments (Arnlund et al., 2014; Cho et al., 2018; Rimmerman et al., 2017). Our instrument configuration allowed us to measure low-angle protein scattering and high-angle solvent scattering simultaneously on the same detector image. To characterize the temperature-dependent behavior of the solvent scattering, we performed static SAXS/WAXS measurements of our CypA samples as a function of temperature (equilibrium, no IR laser), in addition to our time-resolved measurements. We pooled these static, temperature-dependent SAXS/WAXS curves (azimuthally integrated *I(q)* v. *q*) with the time-resolved SAXS/WAXS curves from our T-jump measurements, and performed singular value decomposition (SVD) on a matrix constructed from the set of pooled curves (Figure 2A). Specifically, each column of this matrix represents a scattering curve, with each row of the matrix corresponding to a *q*-bin and the entries in the matrix corresponding to measured scattering intensities. The SVD analysis, which was performed over the *q*=0.07-3.45 region of the scattering curves, identified a signal (a left singular vector) whose prominent features were found in the q-region corresponding to the scattering of bulk water (*q* > 1.0) (Figure 2A). By extracting the entries in the corresponding row of the V matrix (containing the right singular vectors as columns), we could determine how this singular vector contributed to each scattering curve and demonstrate that its contribution was strongly temperature-dependent. Specifically, in the static (no T-jump) scattering curves the contribution of this singular vector increased with temperature, and for T-jump measurements the contribution of this vector to the observed scattering curves is perfectly correlated to the application of the pump laser pulse over sequential laser on-off pairs of X-ray measurements (Figure 2A), providing positive confirmation of a T-jump.

The identification of a temperature-dependent singular vector provided a simple way to measure the magnitude of the laser-induced T-jump. For each of the five static temperatures we explored, we calculated the average value of v_2,n_, the entry in the matrix V that describes contribution of the temperature-dependent singular vector (U_2_) to the nth scattering curve, across 32 individual X-ray scattering curves. We then plotted the average v_2,n_ vs. temperature and fit the data using both linear and quadratic models (Figure 2B). We examined the residuals for the two fits, determined the quadratic fit produced the most appropriate “standard curve” for estimation of the sample temperature from the SVD analysis, and used the resulting second-degree polynomial to estimate the temperature for each scattering curve in our series of time-resolved measurements. We compared the temperatures calculated for neighboring laser on and laser off scattering curves, and found the average T-jump produced by our IR heating pulse to be approximately 10.7°C on average. The SVD analysis also allows us to judge when cooling of the system becomes significant, so that we can identify the maximum pump-probe time delay that is valid for our relaxation analysis (Figure 2A). We observe that v_2,n_ is consistent as a function of pump-probe time delay out to delay times of approximately 562μs, and decreases for longer time delays, implying that significant cooling of the sample takes place in less than 1 millisecond. Consequently, we limited our subsequent analysis to time delays shorter than 562μs. Additionally, we note that following laser T-jump, the solvent reaches a new thermal equilibrium faster than the measurement dead time of our experiment (562ns). This observation is consistent with other work, in which changes to the structure of bulk solvent following laser T-jump have been shown to equilibrate within roughly 200ns (Gruebele et al., 1998).

**Figure 2.**
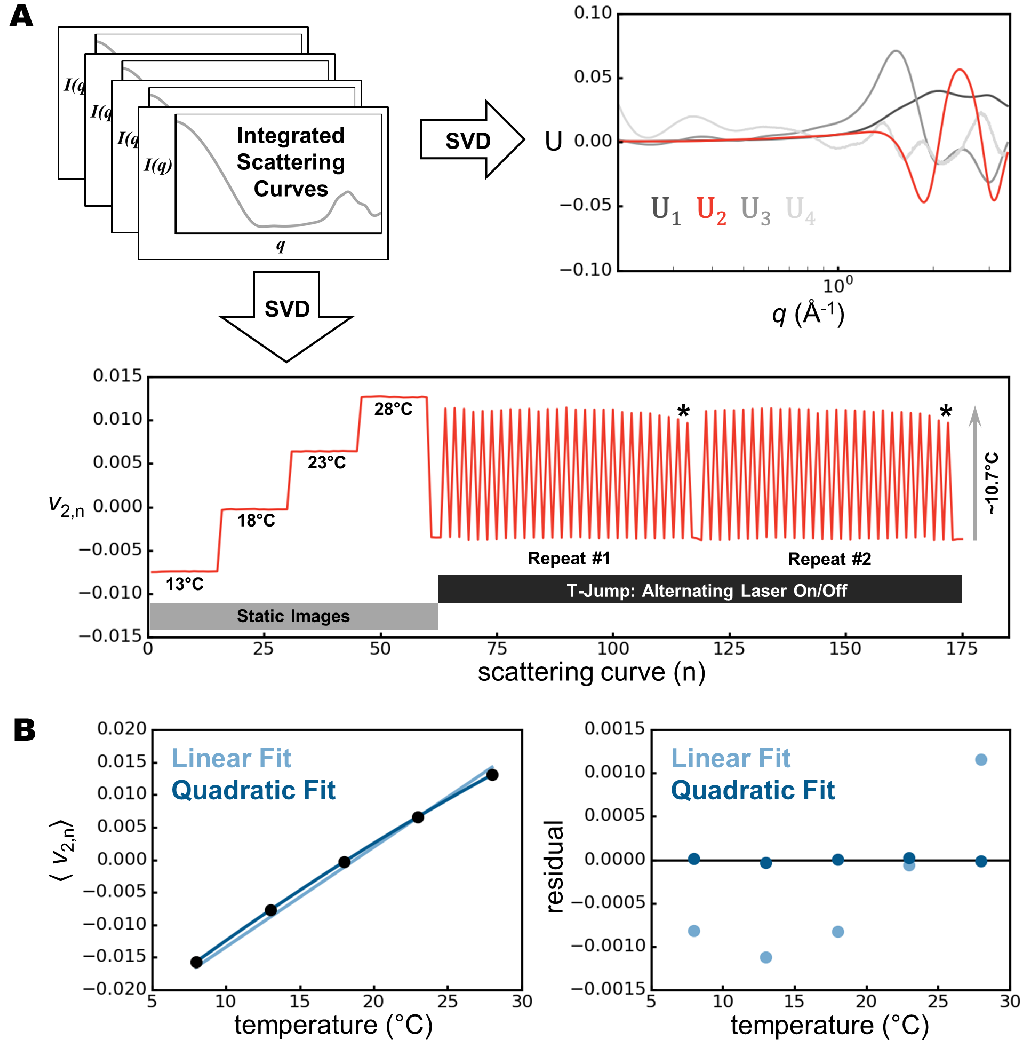
X-ray scattering from bulk water acts as a sensitive thermometer for T-jump experiments. A) Using singular value decomposition (SVD), we can identify a signal whose contribution to each scattering curve is strongly dependent on the temperature. The left singular vectors with the four highest singular values are shown, with the vector corresponding to the temperature-dependent signal (U_2_) colored red. The contribution of this vector (v_2,n_) to each of 175 scattering curves is also shown. This set of 175 scattering curves includes static measurements (no pump laser) at four different temperatures, followed by two repeats of time-resolved T-jump measurements. The T-jump data were collected as laser on-off pairs, and within a single repeat each successive on-off pair was collected with an increasing pump-probe time delay. Cooling is evident at longer pump-probe time delays (denoted by *). B) To calculate the magnitude of the laser-induced T-jump, we used the static data to determine the average value of v_2,n_ as a function of temperature, and fit the data using both linear and quadratic models. Based on the residuals for the two fits, we chose to use the resulting quadratic equation to determine the magnitude of the laser-induced T-jump using the values of v_2,n_ calculated for the time-resolved scattering curves by SVD.

### T-Jump Produces Changes in the X-ray Scattering Profile of CypA

To determine the effect of the T-jump, we initially averaged all data for a given time delay, examined the scattering profiles for differences (Figure 3), and observed a small laser-induced change in the low-*q* region of the scattering profiles. Next, we sought to increase the sensitivity of the experiment by exploiting the structure of the interleaved data collection (Figure 1B). We calculated the “on-off difference” between each set of paired “laser on” and “laser off” scattering profiles. Next, we binned the on-off difference scattering curves according to the associated pump-probe time-delay, performed an iterative chi-squared test to remove outliers (*χ*^2^=1.5), and averaged the calculated differences for all repeat measurements (Figure 3). This subtraction and averaging resulted in accurate measurements of laser on-off difference signals as a function of the pump-probe time delay, and revealed changes at low-*q* (0.03-0.3A^−1^), which we analyze below in the context of the protein’s physical dimensions and overall scattering mass, and high-*q* (1.0-4.2A^−1^), which we used to calibrate the final temperature after laser illumination. The procedure used to determine the sample temperature from the observed X-ray scattering is described in detail above; however, it is worth noting here that the shape of the on-off difference signal at high-*q* is nearly identical to the left singular vector used to monitor the temperature by SVD. The same T-jump measurements were performed on protein samples and on samples consisting of buffer only without protein. After an additional scaling step, average on-off differences for the buffer alone were subtracted from average on-off differences for the buffer with protein, which isolated the signal changes at low-*q* due only to the protein.

**Figure 3.**
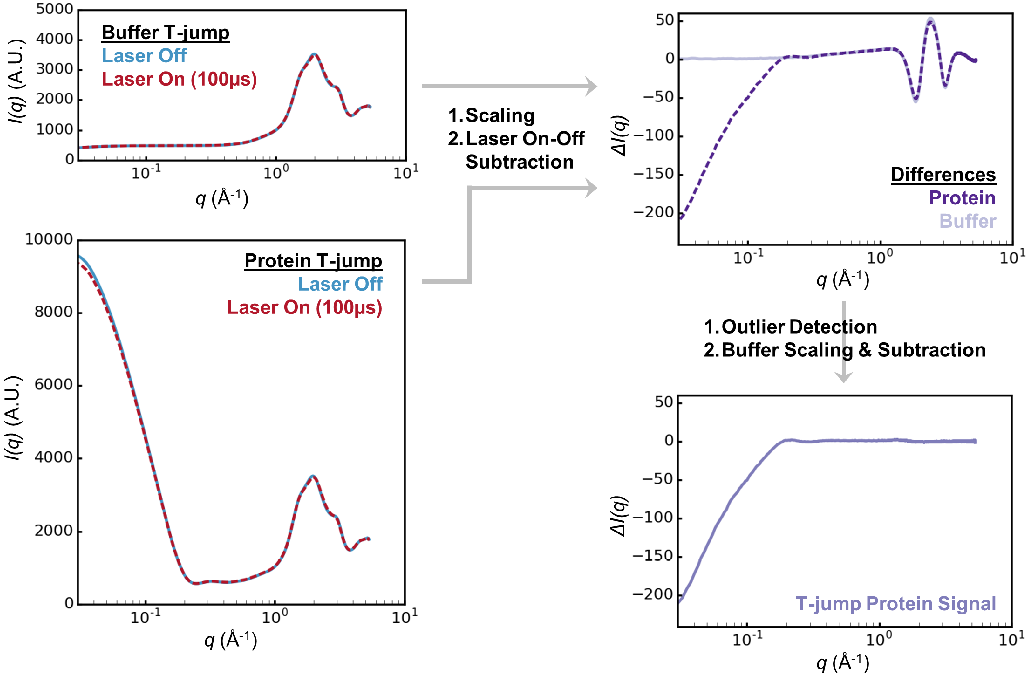
T-jump data processing involves a combination of scaling and subtraction operations that produce time-resolved difference scattering curves. For each laser on-off pair, the recorded scattering curves are scaled to one another and the laser off curve is then subtracted from the laser on curve. This procedure is done independently for samples containing buffer only, and for protein samples. Next, the resulting difference curve for the buffer only sample is scaled to the difference curve obtained for the protein sample, and an additional buffer subtraction is performed to remove the thermal signal from the solvent. The result of this procedure is a difference scattering curve containing signal from the protein molecules only.

### Time-Resolved Changes in Small Angle X-ray Scattering

Comparison of difference scattering curves calculated for 27 time delays revealed a time-dependent change in X-ray scattering by the protein, demonstrating that the modest T-jump we introduced was capable of exciting protein dynamics that could be observed in real time. The difference curves calculated from our data, showing the contribution of the protein to time-resolved changes in the SAXS/WAXS signal, have features in the low-*q* (*q*=0.03-0.20Å^−1^) region (Figure 4A). Qualitatively, the time-resolved on-off differences show that the overall low-angle scattering and extrapolated value of *I(0)* are reduced within the dead time of our experiment (562ns), and then begin to increase slightly over the next few microseconds before decreasing further at longer pump-probe time delays out to 562μs. Changes in low-angle scattering and *I(0)* reflect changes in the overall size and shape of the particles in solution, with a reduction in both observables indicating of a loss of scattering mass and shrinkage of the scattering particles. The observed laser on-off difference in *I(0)* is approximately 3% of the total observed signal, with one-third of that signal change occurring in the time regime that can be resolved by our measurements.

**Figure 4.**
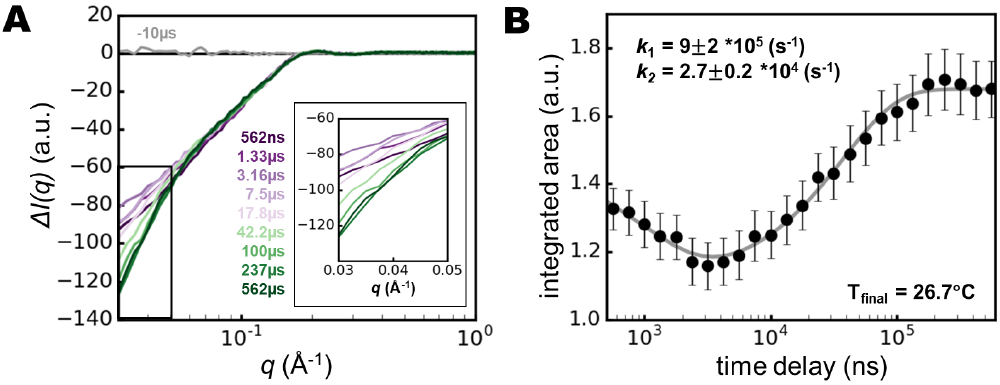
Time-resolved T-jump data allow kinetic modeling of conformational dynamics. A) A series of time-resolved difference X-ray scattering curves is shown for a subset of our T-jump data (10 out of 26 unique time delays). Data at low *q* are plotted on a linear *q* scale in the inset. B) The area under the difference scattering curve in the *q*=0.0275-0.04 region was integrated for all measured pump-probe time delays, and the resulting absolute values are plotted as a function of the pump-probe time delay. The plotted data suggest the existence of multiple relaxation processes, and we used a two-step model of relaxation kinetics to fit the observations (gray line). The rates calculated from the kinetic fit are provided.

The on-off difference for our shortest pump-probe time delay (562ns) is significantly different from 0 at low scattering angles, which suggests the existence of structural changes in the system that are faster than the dead time of our measurements. The physical basis for the fast signal change is likely due to a combination of thermal expansion and change in the amount of ordered solvent surrounding the protein. First, thermal expansion of solvent results in the expulsion of some scattering mass from the X-ray beam path. Based on the volumetric thermal expansion coefficient of water (approximately 0.0003/°C) (Irvine and Duignan, 1985), this effect reduces the overall scattering mass by approximately 0.3% for our T-jumps, which were approximately 11°C. Second, protein thermal expansion coefficients are estimated to be larger than those of liquid water (Frauenfelder et al., 1987; Hiebl and Maksymiw, 1991), so heating of the sample could result in a reduction in scattering contrast between the protein particles and the bulk solvent. Thermal expansion of solvent is well-known to occur within approximately 200ns, and it is reasonable to assume that protein thermal expansion may occur on a similar timescale. The thermal expansion coefficient of CypA is unknown, and rates of protein thermal expansion in general have not been studied explicitly, although measurements of “protein-quake” motions in photoactive systems suggest these effects likely occur within hundreds of picoseconds (Arnlund et al., 2014; Levantino et al., 2015). Kratky plots created from our static and T-jump data suggest a slight increase in protein flexibility without unfolding (**Supplemental Figure 2**), which we interpret to be the result of protein thermal expansion and an overall increase in thermal disorder. This process appears to be faster than the dead time of our measurements, since the effect is temperature-dependent, but not time-dependent over the pump-probe time delays we explored. In addition to thermal expansion effects, the temperature change likely causes some of the ordered solvent around the protein to “melt” into the bulk, which could also lead to a fast decrease in the overall scattering mass and size of the protein particle. The kinetics of these fast processes, while potentially interesting, are invisible to our experiment. Therefore, our subsequent analysis is focused on structural dynamics that occur in the microsecond regime.

Because the main time-resolved signal change was confined to very low-*q*, we wanted to ensure our time-resolved signal was due a change in the protein’s form factor (infinite dilution), and not the structure factor of the protein solution. To test whether changes in the radial distribution function originated from structural changes within the individual protein particles and their associated solvent and not from changes in the relative arrangement of the CypA molecules in solution, we performed static SAXS/WAXS measurements of CypA as a function of both temperature and CypA concentration (**Supplemental Figure 3**). This control allowed us to characterize and correct for the effect of interparticle interactions. Using these data, we calculated the structure factor (*S(q)*) for a 50mg/mL CypA solution at multiple different temperatures, and determined there was no significant difference in the *q*=0.03-0.2 region of the scattering curves, consistent with other work on similarly-sized protein molecules in solution (Bonneté et al., 1999). Next, we plotted the second virial coefficient for CypA as a function of temperature, and noticed that this quantity shows only a very small temperature dependence that cannot account for the observed time-resolved differences. In contrast to our results for CypA, Bonneté, et al. performed similar calculations of second virial coefficients for lysozyme solutions at similar temperatures and buffer conditions, and calculated temperature-dependent changes that were 50-fold larger (or more) than what we determined for CypA (Bonneté et al., 1999). In addition to the direct measurements of interparticle interactions provided by concentration-dependent scattering measurements, we also used Guinier analysis to assess whether the radial distribution function (structure factor) of CypA particles in solution changes significantly upon temperature perturbation. We performed linear fits of *ln[I(q)]* vs. *q*^*2*^ for averaged scattering curves derived from static temperature data and from time-resolved data, and observed that the residuals do not change substantially as a function of either temperature (in static experiments) or time (in time-resolved experiments). Because deviations from the linear Guinier fit are often the result of interparticle interactions, we concluded that the relative consistency of these residuals provides additional evidence that such interactions have a negligible effect on our observations. The Guinier analysis was also used for structural interpretation of the time-resolved signal, which is described in detail below.

Comparison of laser on and off scattering curves revealed scattering differences that were approximately the same in magnitude and direction as differences between static temperature measurements performed on samples equilibrated to temperatures that differ by 10°C (roughly corresponding to the magnitude of the laser-induced T-jump, **Supplemental Figure 4**). The equilibrated signal change that we observe in our time-resolved measurements (*I(0)*_562us_ - *I(0)off*) is similar, but not identical, in magnitude (3.2% of total) and direction to differences calculated using static scattering curves collected at temperatures that approximate the laser on and laser off measurements in our time-resolved experiments (1.9% of total signal). The small discrepancy in the overall signal change induced by a ~10°C temperature change in static versus dynamic experiments could be due to additional relaxation processes which occur on timescales longer than we measure in our experiment (our measurements extended out to 562μs), whereas some motions in CypA have been reported to have millisecond exchange rates (Eisenmesser et al., 2005), or due to systematic errors in comparing static and time-resolved measurements.

**Figure 5.**
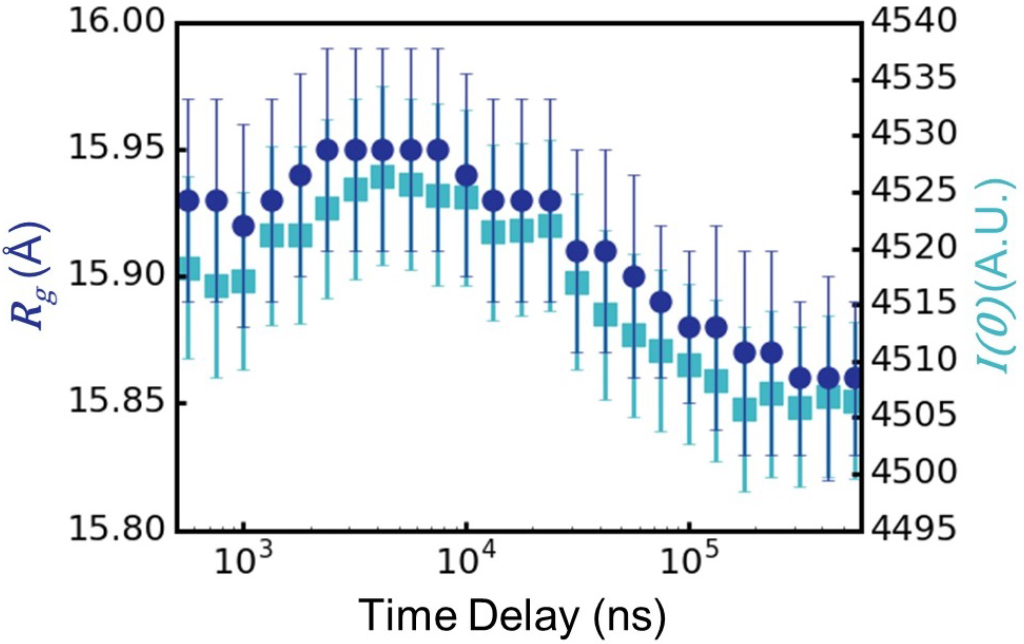
Guinier analysis can be used to estimate changes in physical parameters of the protein particles from the timeresolved scattering data. Consistent with our kinetic analysis, the radius of gyration (*R*_*g*_) of the average CypA particle in solution first increases and then decreases as a function of time following the T-jump. Additionally, the value of *I(0)* extrapolated from the Guinier analysis shows a simultaneous increase and decrease, suggesting that the change in the particle size is coupled to a change in its scattering mass, which is likely due to the acquisition and loss of water molecules from the solvation shell as the protein swells and then shrinks.

### Kinetic Modeling of Structural Dynamics from Time-Resolved Scattering Differences

Our time-resolved measurements of scattering differences allowed us to model the kinetics of global structural changes induced by the T-jump. For kinetic modeling, we integrated the area under each of our time-resolved difference curves in the *q*=0.0275-0.04 region and plotted the absolute value of the area as a function of the associated pump-probe time delay (Figure 4B). Based on the apparent shape of the area vs. time delay plot, we reasoned that a two step kinetic model would be needed to fit the data, since the area first decreases, and then increases, as a function of time delay. We fit the observed data to a two-step model of relaxation kinetics (independent steps) using a non-linear least-squares curve fitting algorithm, and calculated rates of 9×10^5^ s^−1^ ± 2×10^5^ s^−1^ for the fast process (*k*_*1*_) and 2.7×10^4^ s^−1^ ± 0.2×10^4^ s^−1^ for the slow process (*k*_*2*_) at 26.7°C (299.7K). The errors calculated for these rates are the result of propagating measurement standard deviations through radial integration, scaling, on-off subtraction, averaging, buffer subtraction, difference curve integration, and kinetic fitting. It is worth noting that the errors calculated in our analyses are likely to overestimate the true error, as we considered all experimental errors to be random. In contrast, some experimental error is likely systematic, and would instead be removed, rather than propagated, by the subtractive operations employed during data processing.

**Figure 6.**
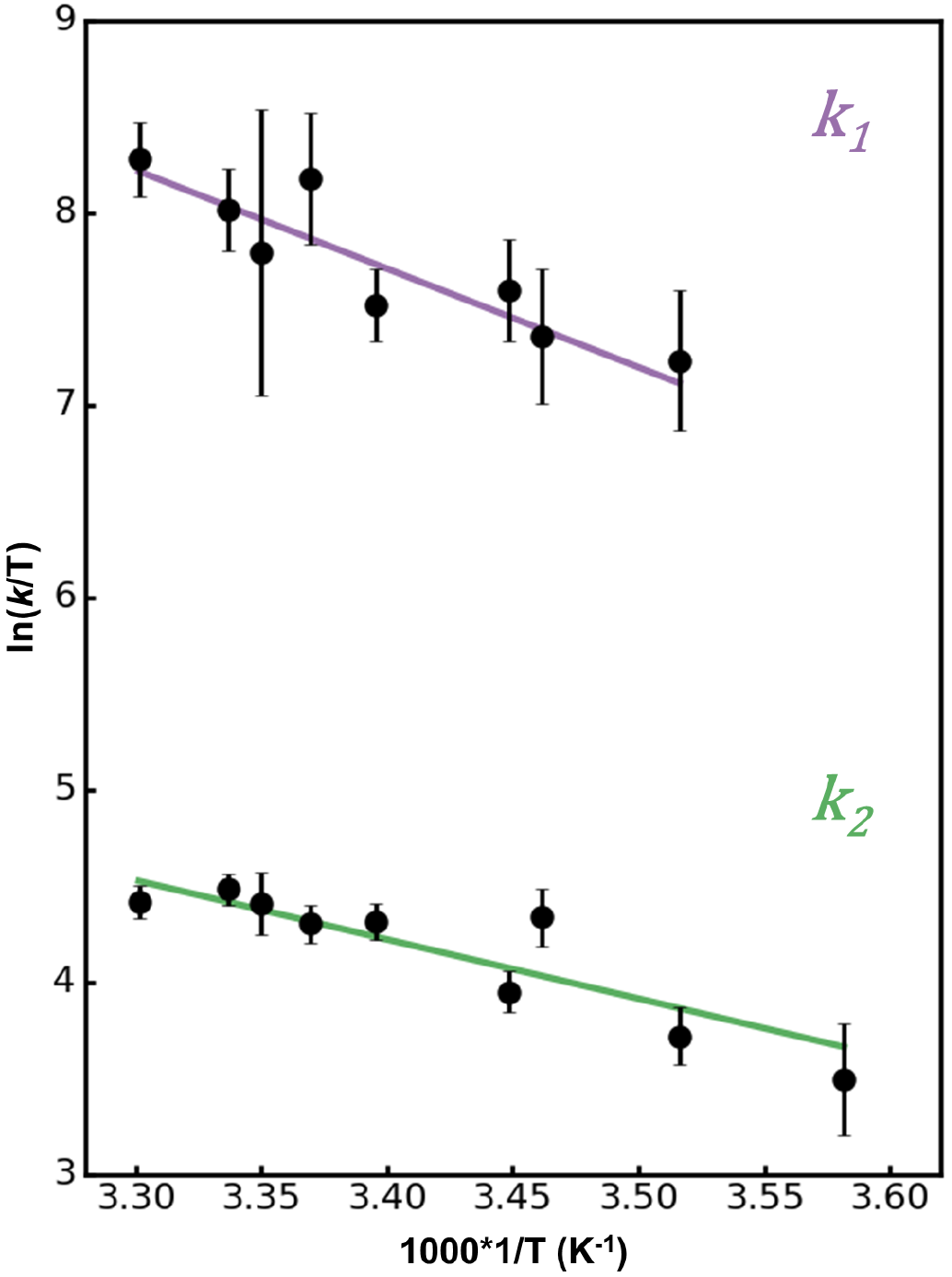
Linear Eyring plots for each of the two relaxation processes observed in our T-jump experiment with CypA. The data are fit using Eq. 1. The linear fit for the fast process (*k*_*1*_) is shown in purple, and the linear fit for the slow process (*k*_*2*_) is shown in green.

In addition to our kinetic analysis of the on-off difference curves, we also used the time-resolved data to generate *I(q)*_*t*_ scattering curves, which we subsequently used for Guinier analysis to determine how the radius-of-gyration (*R*_*g*_) of the CypA particle changes as a function of time following the T-jump (Figure 5). After the T-Jump, the average CypA particle shrinks in the dead time of our experiment; however, within a few microseconds of the T-jump, a fast structural transition (described by *k*_*1*_ in our kinetic analysis) causes the CypA particle to expand subtly. While the increase in the calculated radius of gyration is small relative to the error on the Guinier fit for any single data point, our conclusion that the particle is expanding is supported by multiple time points and kinetic analysis of the integrated area under difference scattering curves. Following this fast increase in *R*_*g*_, a second, slower process (described by *k*_*2*_ in our kinetic analysis) reverses this trend, causing the average CypA particle to shrink again.

Next, to learn more about the conformational transitions in CypA that are excited by the T-jump, we repeated the experiment at multiple different jumped temperatures ranging from 6.2°C to 29.9°C (279.2K to 302.9K). We modeled the kinetics of the SAXS/WAXS signal changes to observe how the relaxation rates changed as a function of temperature. The calculated rates (*k*_*1*_ and *k*_*2*_) for all temperatures are provided in Table 1. We analyzed the temperature-dependence of these rates using the Eyring equation, which provided insight into the thermodynamics of the transition states for the two processes. First, we plotted *ln(k/T)* versus *1/T* (Figure 6), and noted that the relationships appeared to be linear. Therefore, we used the fitted slopes and y-intercepts to calculate the enthalpies and entropies of activation for each of the two processes according to the linearized Eyring equation:

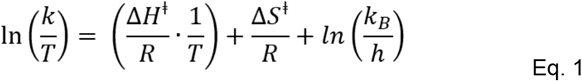

where *R* is the gas constant, *k*_*B*_ is the Boltzmann constant, and *h* is Planck’s constant. The enthalpies of activation (*ΔH*^*‡*^) and entropies of activation (*ΔS*^*‡*^) and their standard deviations are given in Table 2. The fast process (*k*_*1*_) has a large, positive enthalpy of activation, but this is partially offset by a positive entropy of activation. Formation of the transition state during the slow process *k*_*2*_) has a smaller enthalpic cost, but is also entropically disfavored. We note that the lowest temperature measurement was not used in the Eyring analysis of the fast process (*k*_*1*_) because the error on the measured rate was large due to the low magnitude of the overall time-resolved signal changes at this temperature.

**Table 1.**
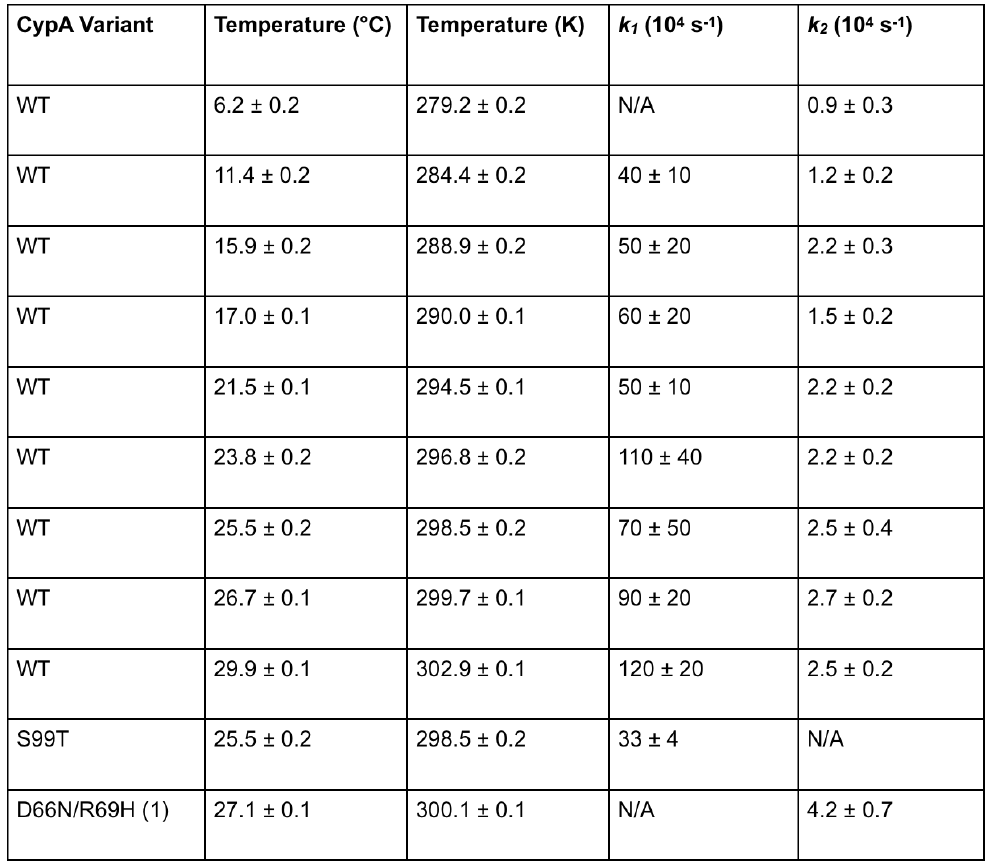
Calculated rates for the fast (k1) and slow (k2) relaxation processes measured from all T-jump experiments reported here. Note that kinetic analyses for the S99T and NH variants were performed at 25.5°C and 27.1°C respectively.

**Table 2.** Enthalpies (ΔH^‡^) and entropies (ΔS^‡^) of activation for the fast (*k*_*1*_) and slow (*k*_*2*_) processes observed for CypA, calculated from Eyring analysis.

| | $\Delta H^\ddagger$ ($10^3$ J mol$^{-1}$) | $\Delta S^\ddagger$ (J mol$^{-1}$ K$^{-1}$) |
| --- | --- | --- |
| Fast Process ($k_1$) | 43 ± 8 | 50 ± 27 |
| Slow Process ($k_2$) | 26 ± 6 | -37 ± 19 |

### CypA mutations with distinct effects on conformational dynamics alter time-resolved signal changes

The time-resolved signal changes that we attributed to WT CypA were observed only at low scattering angles, and therefore the resulting structural information had very limited resolution. To gain a better understanding of the structural transitions excited by the T-jump, we next studied two specific CypA mutants, S99T (in the “core” region, Figure 1B) and NH (D66N/R69H in the “loops”, Figure 1B). The conformational dynamics of these two variants of the enzyme each differ from the wild type in distinct ways: S99T is catalytically impaired due to a loss of rotameric exchange in a key network of residues, whereas NH alters the substrate specificity of CypA by enhancing the dynamics of the surface-exposed loops adjacent to the active site. Importantly, NMR relaxation measurements indicate that S99T perturbs the active site but not the loops (Fraser et al., 2009), and that NH only perturb the loops (Caines et al., 2012).

**Figure 7.**
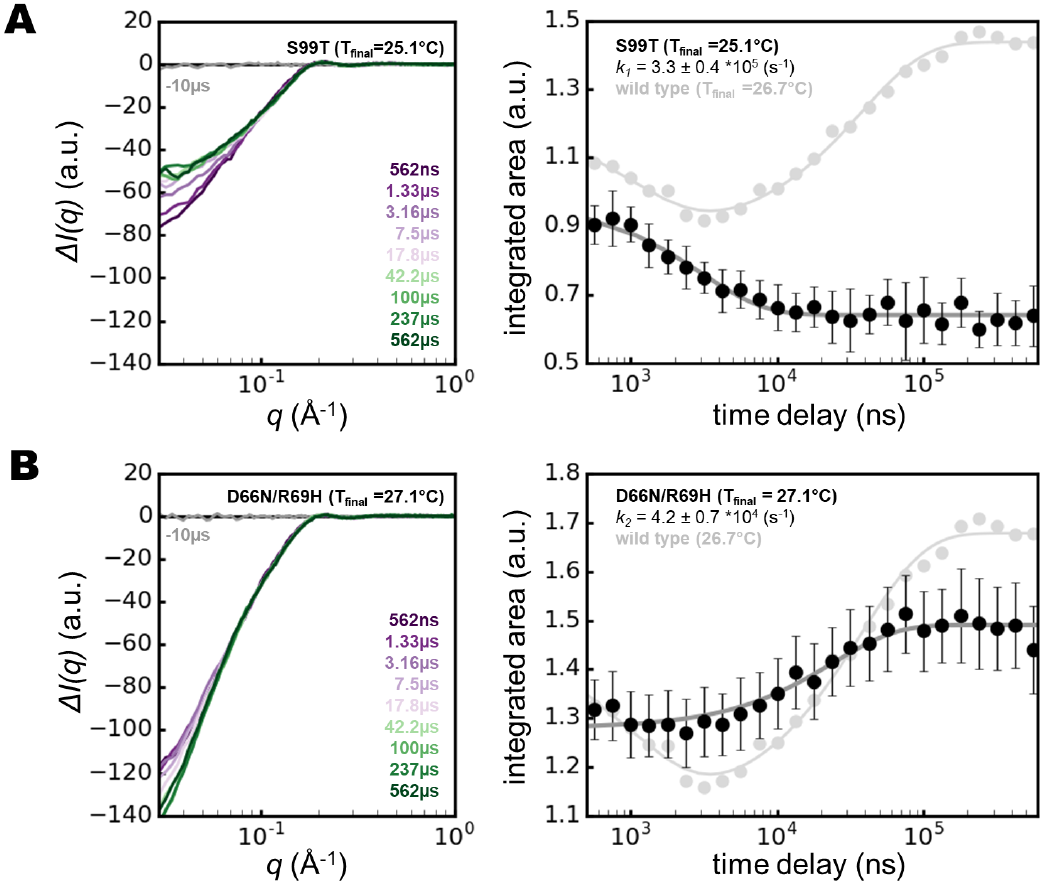
Kinetic analysis of two CypA mutants with distinct effects on the enzyme’s function demonstrate the link between the observed T-jump signal and functional dynamics. The data are presented in the same manner as for the wild type enzyme, shown in Figure 4. In the plots of integrated area versus pump-probe time delay (right panels), the signal observed for the wild type enzyme is shown in light gray for comparison. A) The S99T mutant, which displays defective catalytic function, shows only the fast relaxation process (*k*_*1*_) and lacks the slower process (*k*_*2*_). Note that in the right panel, the gray curve representing the wild type signal is offset by approximately −0.2 units, which accounts for the difference in integrated area due to a beamstop shift during the S99T measurements relative to measurements of other variants. B) The D66N/R69H (NH) mutant, with altered substrate specificity, shows the slow relaxation process (*k*_*2*_) and lacks the faster process (*k*_*1*_).

We observed that both S99T and NH mutants showed time-resolved SAXS signal changes that differed from the wild type enzyme. Both mutants show a fast signal change that occurs within the measurement dead time of the experiment, which is similar to what we observed for the wild type enzyme and consistent with these changes being largely due to temperature-dependent changes to the solvation shell and increased thermal disorder. Beyond the initial fast loss of scattering intensity that was observed for all three CypA variants we studied (WT, S99T, and NH), the evolution of the time-resolved signals for each of the two mutants differ substantially from the wild type and from one another. In the S99T mutant (Figure 7A), we observed only the fast decrease (*k*_*1*_) of the integrated area under the difference curve (*q*=0.03-0.04), and a striking absence of the subsequent increase (**k*_*2*_*) in the integrated area at longer time delays that was observed for the wild type enzyme. In contrast, for NH, the plot of integrated area under the difference curve as a function of time delay (Figure 7B) appears to lack the initial fast decrease (*k*_*1*_), but it does appear to retain the slower signal change (*k*_*2*_) that results in an increase for this quantity at longer time delays. We initially fit the data from both the S99T and NH variants using a two-step relaxation model, as we did for the wild type. We found that for the mutants, the two-step kinetic model yielded at least one rate with a large error. For the S99T mutant, the first step (*k*_*1*_) was well fit but the second step (*k*_*2*_) was poorly fit, while the opposite was true for the NH variant. After visual inspection, we chose to use a single step kinetic model to fit the data for the S99T and NH mutants, and the calculated rates for the two mutants (*k*_*1*_ for S99T and *k*_*2*_ for NH) are also given in Table 1. Plots of the residuals for these fits revealed no structure, suggesting that a single-step kinetic model is sufficient to explain the data for the CypA mutants. In contrast, fitting kinetic data collected for the wild type enzyme using a single-step model results in residuals with exponential character, and a two-step kinetic model is needed to reduce the error in the fit (**Supplemental Figure 5**).

Our measurements of the S99T and NH variants of CypA clearly demonstrated that mutations which are known to impinge on the activity and specificity of the enzyme also perturb the observed time-resolved signal relative to the wild type in our T-jump experiments. Most notably, the slow relaxation process (modeled by *k*_*2*_) is shared only by the catalytically-competent wild type and NH variants, and its absence from the S99T variant suggests that the underlying conformational change is related to the catalytically-coupled motions that are arrested by the S99T mutation. These results indicate that T-jump experiments are capable of exciting and measuring functionally-relevant, intramolecular structural dynamics of proteins, even when the data are limited to relatively low scattering angles.

## DISCUSSION

These results demonstrate the utility of T-jump X-ray scattering experiments for characterizing the intramolecular structural dynamics of proteins. Time-resolved T-jump X-ray scattering experiments have the potential to be a powerful tool for understanding the complex dynamics of protein molecules, such as the model enzyme CypA, which has no intrinsic photoactivity. In our experiments, T-jumps of approximately 10-11°C modified the CypA conformational ensemble, producing a clear, time-dependent change at very low scattering angles. High-angle scattering differences required for atomistic structural interpretation were not observed due to signal-to-noise considerations. We systematically ruled out the possibility that low-angle scattering changes were due to the temperature-dependence of interparticle spacings (quantified by the structure factor, *S(q)*), and subsequent Guinier analysis of time-resolved scattering curves allowed us to track changes in the average radius-of-gyration (*R*_*g*_) of the scattering particles, which include the CypA molecules plus their solvation shells of ordered water molecules. This signal change is comprised of an initial reduction in low-angle scattering that occurs within the measurement dead-time of our experiment, followed by a small increase in low angle scattering that equilibrates within a few microseconds (*k*_*1*_ = 9 ±2 ×10^5^ s^−1^ at 26.7°C), and finally a further reduction in low angle scattering that equilibrates within tens of microseconds (*k*_*2*_ = 2.7±0.2 ×10^4^ s^−1^ at 26.7°C).

This analysis suggests that the scattering mass of CypA (and ordered solvent) first swells and then shrinks after excitation by the T-jump. By performing T-jump experiments over a range of temperatures, we discovered that both the fast and slow processes we observed could be described using Arrhenius kinetics. An Eyring analysis revealed relatively large, positive enthalpies of activation for both processes, consistent with the idea that conformational changes generally require breakage of existing interactions in both the protein and in the solvent, as required by the “solvent slaving” concept. The activation enthalpy for the fast process (*k*_*1*_) is larger, but the overall activation energy is lower because of a favorable activation entropy. The opposite is true for the slower process (*k*_*2*_), which has a smaller overall activation enthalpy, but has a disfavorable activation entropy.

We used the S99T and NH variants of CypA to disentangle the nature of these processes and their associated functions. The S99T mutant is capable of undergoing the fast (*k*_*1*_) expansion process, but does not experience the subsequent slow (*k*_*2*_) shrinkage. NMR and crystallography have shown that this mutation arrests the conformational exchange of the “core” catalytic network of residues in CypA by creating steric hindrance, strongly favouring a minor conformation of the wild type enzyme (Otten et al., 2018). This interpretation suggests that the internal rearrangements are related to the *k*_*2*_ process. However, there is a separation of timescales between the NMR results, which indicate ms dynamics in the “core” region, and the T-jump SAXS results here, which indicate μs dynamics. This discrepancy may reflect coupled processes that are related by a population shuffling mechanism (Smith et al., 2015) and agree with a broad timescale range of side chain dynamics in CypA uncovered by molecular dynamics experiments (Wapeesittipan et al., 2018). In contrast to the S99T mutant, the NH variant lacks the initial fast signal change (*k*_*1*_) in our T-jump experiments, but clearly retains the slow (*k*_*2*_) signal. NMR and crystallographic studies of the D66N/R69H (NH) double mutant demonstrated that it maintains wild type catalytic motions, but enhanced flexibility of a surface loop region adjacent to the active site (Caines et al., 2012). This change is due to breaking of several hydrogen-bonding interactions and leads to changes in substrate specificity. Therefore, we hypothesize that the loop motions are responsible for the fast (*k*_*1*_) signal in WT CypA and S99T, where these motions are known to be unperturbed by NMR (Fraser et al., 2009). However, NMR experiments with NH have shown that the loop motions still occur, but at an increased rate that renders them invisible to our experiments. Furthermore, the assignment of these motions by the mutational analysis is consistent with the *R*_*g*_ changes observed experimentally during each process. X-ray crystal structures indicate that the minor conformational state of the catalytic network and associated solvent are smaller than the major state and its associated solvent. Using a room temperature X-ray crystal structure of wild type CypA (PDB: 3K0N), we calculated the radius-of-gyration of the enzyme with the core catalytic network (Arg55, Met61, Ser99, and Phe113) in both the major and minor conformational states, and found that the predicted Rg of the minor state is 0.07Å smaller than the major state (14.09Å vs. 14.16Å). Additionally the increase in the average Rg during the faster process (*k*_*1*_) is consistent with the loops sampling an expanded conformational ensemble, as indicated by recent exact-NOE NMR ensembles (Chi et al., 2015). Our kinetic modelling of the WT and mutant data suggest that two uncoupled dynamic modes are observed with different kinetics, each of which is individually perturbed by different mutations.

Time-resolved X-ray structural measurements are critical for decoupling the experimental signatures of conformational changes that can become convoluted by the spatial and temporal averaging that is inherent to traditional X-ray experiments. If one were to assess traditional, static SAXS data for CypA, one would find that increasing the temperature of the sample results in a decrease in the average particle size at equilibrium. These static measurements as a function of equilibrium temperature fail to capture that the temperature change actually perturbs two distinct protein motions, which have the opposite effect on the enzyme’s global structural characteristics. This information can only be obtained through a time-resolved experiment, which is able to separate the effects of these two motions because they have substantially different rates. The ability to dissect individual conformational motions and measure their rates using time-resolved X-ray measurements is important for understanding processes involving complex protein dynamics. Many of these dynamic processes, including allostery (Colombo et al., 1992; Kim et al., 2016; Royer et al., 1996; Salvay et al., 2003) and enzyme catalysis (Decaneto et al., 2017; Fenwick et al., 2018; Grossman et al., 2011; Guha et al., 2005; Leidner et al., 2018), involve extensive reorganization of interactions between the protein and its ordered solvation shell, which are key contributors to the energetics that govern protein motions (Caro et al., 2017; Conti Nibali et al., 2014; Dahanayake and Mitchell-Koch, 2018; Fenimore et al., 2002; Frauenfelder et al., 2007; Gavrilov et al., 2017; Wand and Sharp, 2018). Because X-ray solution scattering experiments report on the structure of a protein and the ordered solvent molecules that constitute its solvation shell (Henriques et al., 2018; Hub, 2018; Svergun et al., 1998; Virtanen et al., 2011), the widespread application of time-resolved SAXS/WAXS experiments will enhance our understanding of how protein motions are driven by solvent dynamics, especially when they can be combined with molecular dynamics simulations to provide atomic scale insight into the underlying structural changes (Arnlund et al., 2014; Berntsson et al., 2017; Brinkmann and Hub, 2016; Takala et al., 2014). In order for these experiments to enter the mainstream of structural biology, it has become necessary to create perturbations that can be applied universally, to any protein of interest, and our results establish that T-jump can be used as a general perturbation method to excite functional intramolecular protein motions for time-resolved X-ray structural measurements. Looking forward, T-jumps can be paired with other perturbations, such as mutations and ligand binding, to answer important questions about how disease alleles or drug molecules impinge on protein dynamics.

## METHODS

### Sample Preparation

CypA samples were prepared as described previously. Briefly, the recombinant protein was expressed in *E. coli* bL21(DE3) cells and purified by liquid chromatography. Cells were lysed by sonication at pH=6.5, the lysate was clarified by highspeed centrifugation, and CypA was captured from the clarified lysate using a HiTrap-SP cation-exchange column (GE Healthcare). The protein was eluted using a sodium chloride gradient, and fractions containing CypA were pooled, and the pH was shifted to 7.5. The resulting solution was applied to a HiTrap-Q anion exchange column (GE Healthcare), and CypA was collected in the column flowthrough. Finally, a polishing step was performed using a Superdex-75 gel filtration column (GE Healthcare). The protein was concentrated to 50mg/mL in buffer containing 20mM HEPES (4-(2-hydroxyethyl)-1 -piperazine-ethanesulfonic acid) buffer at pH=7.5, 50mM sodium chloride, and 0.5mM TCEP (tris-hydroxyethylphosphine). CypA mutants (S99T and NH) were prepared following the same protocol used for the wild type protein. For all X-ray measurements performed on buffer only without protein, the buffer was taken from the concentrator filtrate.

### T-Jump SAXS/WAXS Data Collection and Processing

Time-resolved SAXS/WAXS measurements of CypA were performed on the BioCARS beamline at the Advanced Photon Source, while the storage ring was operating in hybrid mode. Temperature-jump data were acquired using the pump-probe method, as described recently by Cho. et al. (Cho et al., 2018). Fast temperature-jump was performed on a CypA solution (50mg/mL) in a silica capillary using an Opolette 355 II (HE) optical parametric oscillator (OPOTEK), which produced a 7ns laser pulse with a peak wavelength of 1443nm. The pump laser energy was approximately 1mJ per pulse, and the beam was focused to an elliptical spot with dimensions of 400pm by 60pm, yielding a photon fluence of ~50mJ/mm^2^ at the sample, which heated a 50mg/mL CypA solution in a capillary. A suitably delayed X-ray pulse of 494ns duration (eight septuplets in APS hybrid mode) with a peak X-ray energy of 12keV and 3% energy bandwidth (pink beam, **Supplemental Figure 1**), was used to probe the sample following the introduction of the T-jump, and the X-ray scattering was recorded using a Rayonix MX340-HS CCD detector. In our experiments, the temporal resolution is limited to approximately 500ns by the duration of the X-ray pulse, which is substantially longer than the duration of the IR pulse. To speed data acquisition, we utilized a sample holder and data collection scheme recently reported by Cho, et al, (Cho et al., 2018) which combined fast translation along the capillary axis with slow sample circulation via a peristaltic pump. The fast translation of the capillary allowed us to rapidly accumulate X-ray scattering from 41 pump-probe measurements on a single detector image by translating the capillary to a fresh position between each pump-probe pair. The slow circulation of the sample allowed us to replenish the protein solution and limit the extent of X-ray radiation damage by spreading the X-ray dose over a relatively large volume during long data collection runs. Data were collected as pairs of alternating “laser-on” and “laser-off” X-ray images. The pump-probe time delay was systematically increased with each successive on/off pair of images. We measured pump-probe time delays spanning three logarithmic decades from 562ns to 1ms, at a time density of eight points per decade. A total of 50 replicate X-ray images were collected for each pump-probe time delay. It is important to note that time-resolved X-ray measurements referred to herein as “laser off,” were followed (10μs) by application of an IR pulse to the sample, as described by Cho, et al. (Cho et al., 2018), which prevented the introduction of a temperature offset created by incomplete cooling in between “laser on” and “laser off” measurements. A temperature controller integrated into the sample holder allowed us to initiate the T-jump from different starting temperatures, and also allowed us to collect static temperature data. Static temperature images were collected in a manner similar to the time-resolved images, but without application of the pump laser pulse. Data collection protocols were identical for protein and buffer samples.

After acquiring the data we applied polarization, geometry, and detector non-uniformity corrections to the 2D X-ray images. The scattering intensities (photons/pixel) were binned and averaged as a function of the scattering vector magnitude (q), yielding isotropic scattering curves (*I(q)* vs. *q*; *q* = *4πsin(θ)*/λ, where *2θ* is the scattering angle and *λ* is the X-ray wavelength) (Cho et al., 2018). Next, for each data collection run, we carried out outlier detection by performing singular value decomposition (SVD) on a matrix constructed from our integrated scattering curves. In this SVD, the left singular vector with the largest singular value represents the global average of all the scattering curves used to construct the input matrix. We analyzed the right singular vectors from the SVD to determine which images were irregular. Specifically, we calculated the mean value of v_1,n_, the entry in the matrix V that describes contribution of the right singular vector with the largest singular value (U_1_) to the n^th^ scattering curve, across all the input X-ray scattering curves. Then, if the value of v_1,n_ for any specific scattering was more than 2.5 standard deviations above or below the mean, that image was discarded. Our outlier detection procedure is implemented in a Python script called “SVD_Quarantine.py.” By inspecting the results of the SVD, we decided to remove the first 5 repeats from each data set, as well as some additional outliers that failed our statistical test. The same averaging and outlier detection method was used for both static and time-resolved measurements.

### Scaling of X-ray Scattering Curves

All scaling of X-ray scattering curves was performed using an algebraic (least-squares) procedure. To determine the scale factor, *A*, which can be applied to a scattering curve *I(q)*_*a*_ in order to scale it to a second scattering curve *I(q)*_*b*_, we used the following equation:

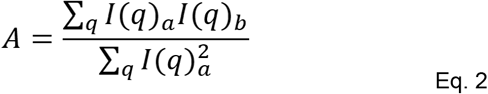

Although we used the equation above for scaling throughout our analysis, the *q*-range to which it was applied varied depending on the context, and details are provided below.

### X-ray Thermometry

Following the initial data processing steps described above, we used singular value decomposition (SVD) to determine the magnitude of the T-jump introduced by the IR laser pulse. We pooled static, temperature-dependent SAXS/WAXS curves (azimuthally integrated *I(q)* v. *q*) with the time-resolved SAXS/ WAXS curves from time-resolved measurements, scaled them to a common reference over the *q*=0.025-5.2 region, and performed SVD on a matrix built from these scaled curves. In this matrix, each column represents a single scattering curve, with the rows of the matrix corresponding to *q*-bins and the entries in the matrix consisting of azimuthally-averaged scattering intensities. The SVD analysis was performed using only the *q*=0.07-3.45 region of the scattering curves. As described in the Results section, the SVD identified a left singular vector whose contribution to the overall scattering signal was highly temperature dependent. This was the left singular vector with the second largest singular value (U_2_). For each of the five temperatures used for static data collection, we calculated the average value of v_2,n_, which is the entry in the matrix V describing the contribution of the temperature-dependent singular vector (U_2_) to the n^th^ scattering curve. We then plotted the average v_2,n_ vs. temperature and ultimately fit this data using a quadratic model. Finally, we used the resulting second-degree polynomial and the values v_2,n_ for each time-resolved scattering curve to estimate the temperature for each T-jump measurement. By comparison of neighboring laser on and laser off scattering curves, we determined that the average T-jump was 10.7°C. Our thermometry procedure is implemented in a Python script called “thermometry_timepoints.py.”

### Data reduction: On-Off Subtraction, Repeat Averaging, and Buffer Subtraction

We implemented a data reduction procedure that operated on the integrated scattering curves generated using our data collection protocol and produced several outputs that were subsequently used for our kinetic and structural analyses. This procedure, implemented in a Python script called “reduce_data.py,” took advantage of paired laser on/off measurements, redundant measurements of each pump-probe time-delay, and parallel T-jump experiments for samples containing protein and samples consisting of buffer only. The input for this script was essentially two data sets. The first, was the series of time-resolved scattering curves measured from a sample containing protein and consisting of paired laser on/off measurements with multiple replicate measurements of each pump-probe time delay (see above). The second was a similar data set, only collected from a sample containing buffer only and no protein. All of the input scattering curves were scaled to a common reference over the *q*=0.025-4.28Å^−1^ range, and “laser off” curves were subtracted from their associated “laser on” curves to create a difference scattering curve (*ΔI(q)*) for each laser on/off pair. Next, all replicate difference curves (i.e. same sample and time delay) were grouped together, an iterative chi-squared test was performed (using a cutoff of *χ*^*2*^=1.5), and the average difference curve was calculated for each pump-probe time delay in the series. For each time delay, the difference signal for the buffer only sample was scaled to the difference signal for the sample containing protein over the *q*=1.5-3.6A-1 range, and then the buffer signal was subtracted from the protein signal to isolate the difference signal due only to the protein. Additionally, this script took all of the “laser off” scattering curves, performed an iterative chi-squared test (cutoff of *X*=15), and calculated their average. As was done for the difference curves, the average “laser off” scattering curve for buffer only was subtracted from the average “laser off” scattering curve for the protein sample after an additional scaling step (*q*=1.5-3.6A-1 range). The output of this data reduction procedure was a single scattering curve (*I(q)* vs. *q*) for the “laser off” state, and a difference scattering curve (*ΔI(q)* vs. *q*) for each pump-probe time delay. All output data were corrected for the contribution of the buffer, and errors were propagated from the initial measurement standard deviations.

### Kinetic Analysis

The averaged difference curves produced by our data reduction procedure were used for kinetic analysis of the time-resolved signal changes, which was implemented in a Python script called “difference_dat_kinetics.py.” For each time delay, this script integrated the area under the difference curve over the *q*=0.0275-0.04 range (*q*=0.03-0.04 for the S99T variant, due to beamstop modifications), then fit the resulting data (integrated area vs. time) to calculate relaxation rates using non-linear least-squares curve fitting. We used the following equations, for single-step kinetic fits:

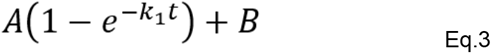

And for two-step kinetic fits:

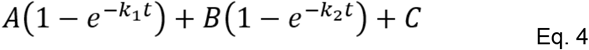

The output of this analysis was a relaxation rate, or two rates, with standard deviations calculated using the covariance matrix from the curve fitting procedure. In cases where we performed T-jumps at multiple temperatures, we used the observed rates and their standard deviations to perform an Eyring analysis by fitting Eq. 1 (above) using a least-squares method to determine the enthalpy and entropy of activation, and their standard deviations (again, using the covariance matrix). We implemented the Eyring analysis in a Python script called “eyring_fit.py.”

### Creation of High-Quality Time-Resolved Scattering Curves for Structural Analysis

To produce high-quality scattering curves that could be used for real space interpretation of the time-resolved X-ray scattering, we took the following steps. We used all of the “laser off” scattering curves from our on-off paired time-resolved measurements to create a single average curve. Then, for each of the time-delays reported, we added the average on-off difference (see above) to this average “laser off” scattering curve:

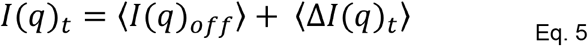

Next, we utilized static scattering measurements, as a function of both concentration and temperature, to characterize the effect of intermolecular interactions on the observed X-ray scattering and to calculate structure factors (*S(q)*) for our 50mg/mL CypA solutions at temperatures spanning a range relevant to our T-jump experiments. We calculated structure factors (and second virial coefficients) following the methods described by Bonnette, et al. (Bonneté et al., 1999). The scattering curves derived from summing the average “laser off” signal and the time-resolved differences were then divided by the calculated structure factors to correct for intermolecular interactions and extrapolate our measurements to infinite dilution. Because we discovered that the effect of intermolecular interactions were not temperature dependent, we did not need to model the time-dependence of structure factors for our protein solutions following the T-jump, and the structure factor calculated at 13°C was used for the infinite dilution extrapolation. The calculation of structure factors and the creation of the high-quality, corrected *I(q)* curves were implemented in a pair of Python scripts called “packing_calc.py” and “packing_correction.py,” respectively.

### Guinier Analysis and Calculation of *R*_*g*_

In order to calculate radii-of-gyration (*R*_*g*_) and to extrapolate the value of *I(0)* from scattering curves, we used the linear Guinier approximation:

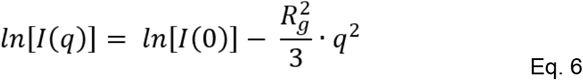

The calculation was implemented in a Python script called “Rg_and_i0.py.”

## Public Availability of Resources

All Python scripts used for analysis of integrated X-ray scattering curves are publicly available on GitHub (https://doi.org/10.5281/zenodo.1493500). Kinetic fits depend on the relax.py tool, which is available separately on GitHub (https://doi.org/10.5281/zenodo.1493508). We are working to make raw data (X-ray scattering images from CCD detector) available via SBGrid following peer review.

## Supporting information

## Acknowledgements

We thank R. Ranganathan, J. Holton and D. Elnatan for helpful discussions, and the staff at the BIoCARS beamline at the Advanced Photon Source (I. Kosheleva, R. Henning, and V. Srajer) for their assistance. This work was supported by: NSF (STC-1231306), NIH (GM123159, GM124149), a Packard Fellowship from the David and Lucile Packard Foundation, UC Office of the President Laboratory Fees Research Program LFR-17-476732 to JSF; the Intramural Research Program of the National Institute of Diabetes and Digestive and Kidney Diseases to PA; a Ruth L. Kirschstein National Research Service Award (F32 HL129989) to MCT. Use of the Advanced Photon Source was supported by the U.S. Department of Energy, Basic Energy Sciences, Office of Science, under contract No. DE-AC02-06CH11357. Use of the BioCARS Sector 14 was also supported by the National Institutes of Health, National Institute of General Medical Sciences Grant R24GM111072. The time-resolved setup at Sector 14 was funded in part through a collaboration with Philip Anfinrud (NIH/NIDDK).

