## Supplementary material for "Temperature-Jump Solution X-ray Scattering Reveals Distinct Motions in a Dynamic Enzyme"

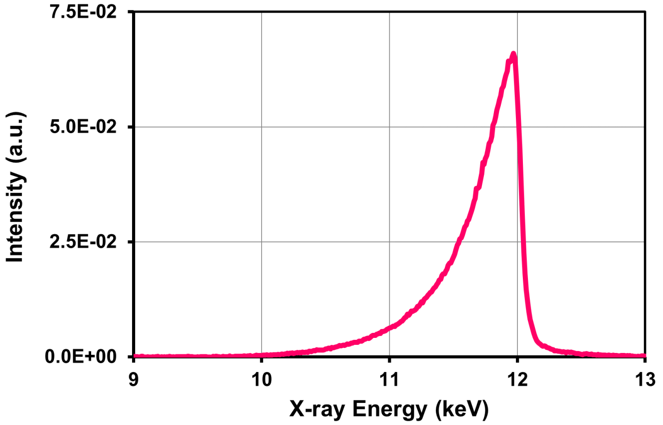


**Figure S1.** Typical X-ray energy spectrum of the pink beam (3% energy bandwidth) used for the reported SAXS/WAXS measurements.

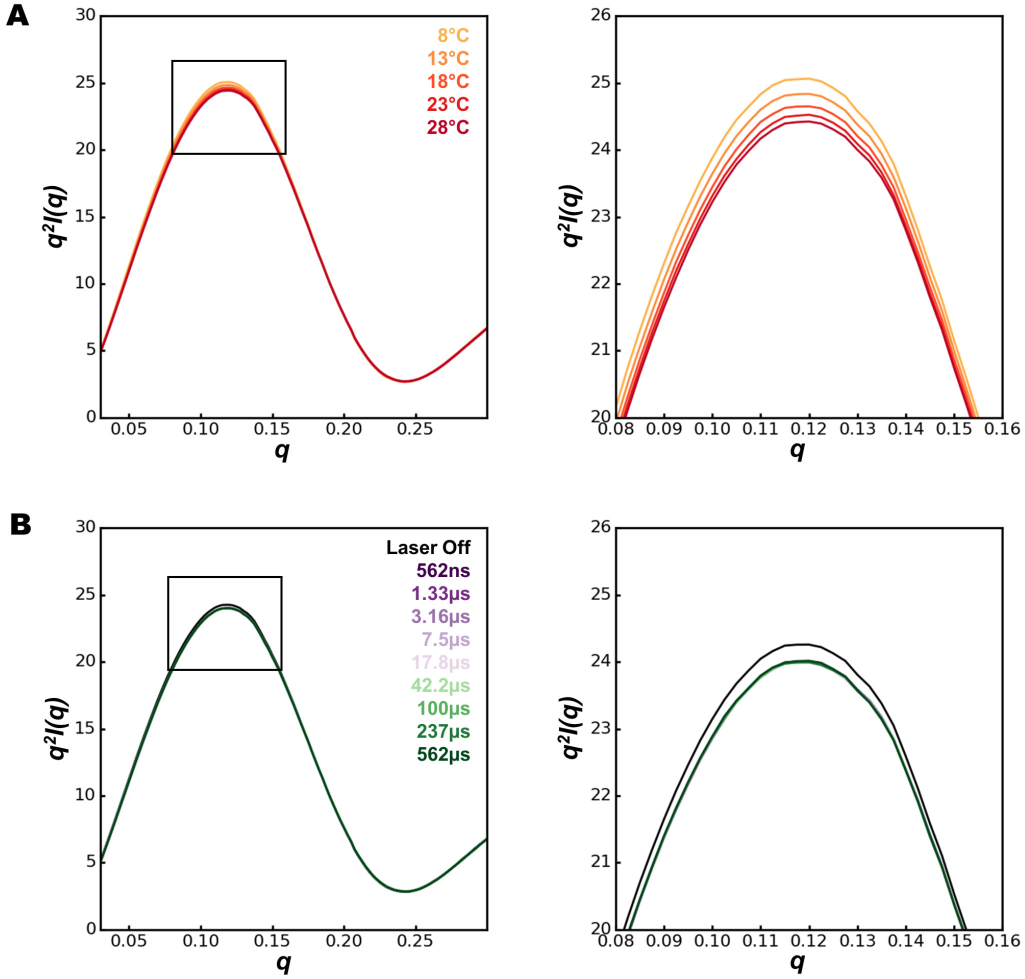


**Figure S2.** Kratky plots for CypA reveal a small thermal disorder effect without protein unfolding. A) Kratky plots calculated as a function of static temperature, from 8°C to 28°C. The right panel shows an expanded view of the boxed region. B) Kratky plots calculated as a function of time delay for time-resolved T-jump data (T-jump from approximately 15°C to 26°C). Again, the right panel shows an expanded view of the boxed region. All time delays show a similar difference relative to the “laser off” state, indicating that the underlying structural change is faster than the measurement dead time of our experiment.

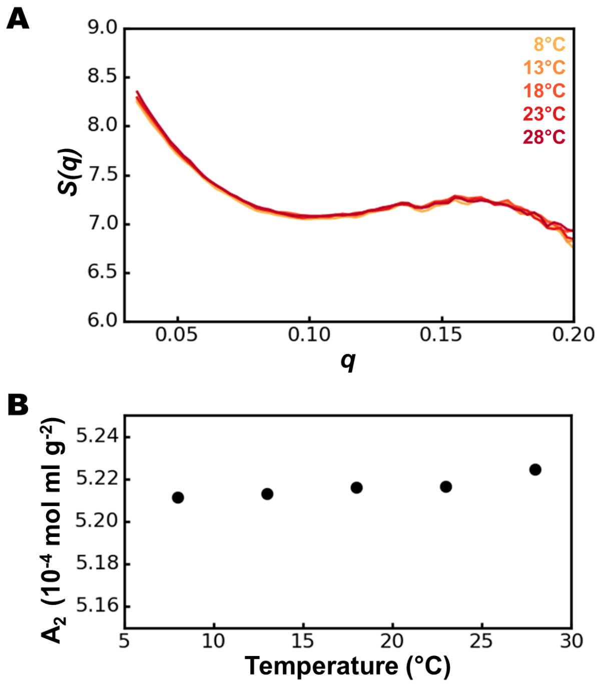


**Figure S3.** Intermolecular interactions are not temperature dependent for CypA solutions. A) Structure factors (S(q)) calculated for 50mg/mL CypA solutions (wild type) at temperatures ranging from 8-28°C. B) Second virial coefficients (A2) calculated for 50mg/mL CypA solutions (wild type) at temperatures ranging from 8-28°C.

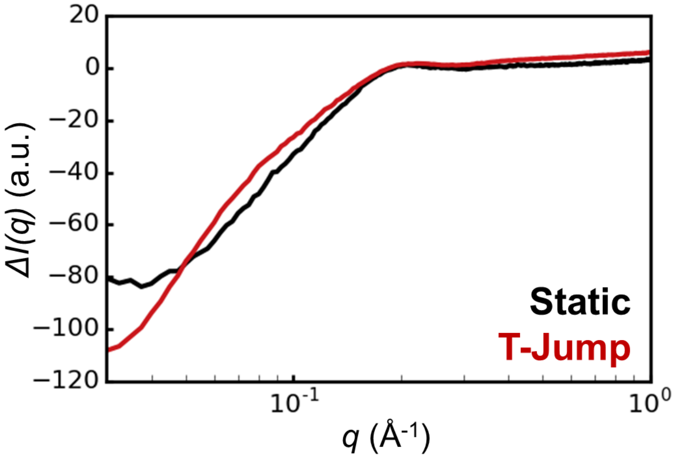


**Figure S4.** Comparison of static scattering differences between CypA solutions at 13°C and 23°C (black curve), and time resolved differences (100μs-laser off) for a T-jump spanning a temperature range of approximately 15°C and 26°C (red curve).

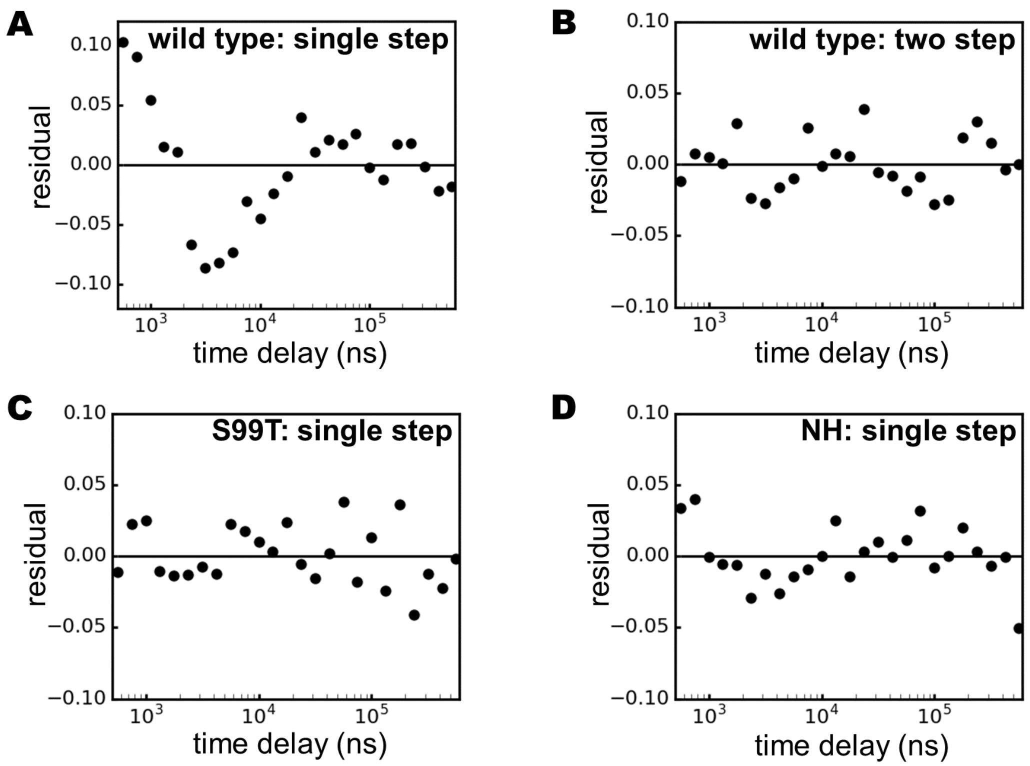


**Figure S5.** Residuals for kinetic fits of T-jump SAXS/WAXS data for CypA variants. A) Wild type, single-step relaxation. B) Wild type, two-step relaxation. C) S99T mutant, single-step relaxation. D) NH double mutant, single-step relaxation.
